# PACT - Prediction of Amyloid Cross-interaction by Threading

**DOI:** 10.1101/2022.07.07.499150

**Authors:** Jakub W. Wojciechowski, Witold Szczurek, Natalia Szulc, Monika Szefczyk, Malgorzata Kotulska

## Abstract

Amyloids are protein aggregates usually associated with their contribution to several diseases e.g., Alzheimer’s and Parkinson’s. However, they are also beneficially utilized by many organisms in physiological roles, such as microbial biofilm formation or hormone storage. Recent studies showed that an amyloid aggregate can affect aggregation of another protein. Such cross-interactions may be crucial for understanding the comorbidity of amyloid diseases or the influence of microbial amyloids on human amyloidogenic proteins. However, due to demanding experiments, understanding of interaction phenomena is still limited. Moreover, no dedicated computational method to predict potential amyloid interactions has been available until now. Here, we present PACT - a computational method for prediction of amyloid cross-interactions. The method is based on modeling a heterogenous fibril formed by two amyloidogenic peptides. The stability of the resulting structure is assessed using a statistical potential that approximates energetic stability of a model. Importantly, the method can work with long protein fragments and, as a purely physicochemical approach, it relies very little on training data. PACT was evaluated on data collected in the AmyloGraph database and it achieved high values of AUC (0.88) and F1 (0.82). The new method opens the possibility of high throughput studies of amyloid interactions. We used PACT to study interactions of CsgA, a bacterial biofilm protein from several bacterial species inhabiting human intestines, and human Alpha-synuclein protein which is involved in the onset of Parkinson’s disease. We show that the method correctly predicted the interactions, performing experimental validation, and highlighted the importance of specific regions in both proteins.

The tool is available as a web server at: https://pact.e-science.pl/pact/. The local version can be downloaded from: https://github.com/KubaWojciechowski/PACT

## Introduction

Pathological misfolding and aggregation of proteins is a hallmark of a number of devastating disorders, including major public health challenges like Alzheimer’s and Parkinson’s diseases^1,2^, type II diabetes^3,4^, as well as some cancers^5^. These diseases not only share a similar molecular mechanism, but they also often co-occur in the same patients. Among others, comorbidities were observed between Alzheimer’s disease and type II diabetes^6,7^ and Alzheimer’s and Parkinson’s diseases^8^. One of the possible explanations of this phenomenon could be related to amyloid cross-interactions. Amyloids are insoluble protein aggregates characterized by exceptional stability due to the tight packing of monomers, resulting in characteristic pattern in X-ray diffraction experiments.^9^. Despite significant structural similarities shared by all amyloids, their sequences are surprisingly diverse and have little homology^10^. On the other hand, sometimes very similar sequences can result in distinctive structures^11^. Numerous, both experimental and computational studies, explored mechanisms of amyloid aggregation and their roles in neurodegenerative disorders, including the pivotal role of oligomers formed at early stages of the aggregation process^12^. More recent studies have shown that in some cases presence of amyloid aggregates can affect the aggregation rate of other proteins^13^. Later, it was observed that interacting proteins can form heterogeneous fibers consisting of molecules of both interaction partners. Hypothetical structural mechanisms of the cross-seeding, depending on the nature of interactors, are proposed in^14^. Aggregation and co-aggregations, observed in the phenomenon of cross-talk, is affected by environmental or experimental conditions. In case of the aggregation enhanced by interaction with another amyloid at conditions hampering the aggregation, the cross-seeding presumably helps to overcome an energy barrier required for fibrillation (e.g., as observed in BSA protein in the presence of HEWL^15^)

The cross-interactions were identified between numerous proteins, including those involved in type II diabetes and neurodegenerative diseases. For example, interactions between Alpha synuclein and human Islet Amyloid Polypeptide (hIAPP)^16^. This shed new light on potentially new aspects regarding the origin of comorbidity of these disorders^17^. A similar mechanism was found to enhance the virulence HIV virus by increasing its adhesion to host cells^18^. Despite the importance of this process, its mechanisms are still poorly understood, although it was shown that polymorphism of an amyloid structure may play a certain role in the aggregation processes^19,20^. The lacking understanding of this process can be attributed to a limited number of experimental data. As a result, interactions of only a few well-described proteins, such as Amyloid-beta (Abeta), islet amyloid polypeptide, or Alpha-synuclein, have been very extensively studied and they contributed to a majority of data. This may introduce a bias in available data. Despite the difficulties, it was shown that proteins with similar sequences are more likely to interact, however, many counterexamples were also shown^17^. The studies highlight the importance of the structural compatibility of amyloid cores.

The main limitation regarding experimental studies of amyloid aggregation and their interactions is that they require expensive and time consuming methods. In practice, biochemical assays based on the binding of Congo Red^21^, Thioflavin T^22^, and infrared spectroscopy are frequently used. Especially the last method is widely applied, due to its simplicity and efficiency^23^. Another approach involves direct observation of fibers using high resolution imaging techniques, such as electron microscopy^24^ and atomic force microscopy^25^. Finally, the advancements in NMR spectroscopy made it an important tool for studying aggregation at the molecular levels^26^. Since different methods rely on different approaches, their results might differ in some cases. More importantly, all of them are expensive and time consuming. Experiments are hampered by difficulties in handling amyloids, including their low solubility, rapid aggregation, and need for their high-purity^27^. Currently, the use of experimental methods for the identification of all amyloids in genome wide studies would be impossible. To address this problem several computational methods have been proposed based on different approaches (reviewed in^28^ and^29^), starting from structural modeling^30^, statistical analysis of the sequence including FoldAmyloid^31^ and FishAmyloid^32^, physicochemical models like PASTA 2.0^33^, machine learning techniques such as APPNN^34^ and AmyloGram^35^. Furthermore, there are methods combining both approaches such as PATH^36^ and Cordax^37^. Finally, some methods, like Aggrescan 3D^38^, utilize information about protein structure. It was also shown that bioinformatics techniques are quite robust and capable of identifying even some misannotated data despite being trained on them^39^. Unfortunately, none of these methods can predict amyloid cross-interactions.

Here, we present a new computational method PACT (Prediction of Amyloid Cross-interaction by Threading) designed for the identification of potentially interacting amyloid pairs. The method is based on the molecular threading applied to the potential complex of interacting amyloids and the assessment of the stability of obtained molecular models.

## Results

The main assumption of the method is that interactions between amyloidogenic fragments that cross-interact, threaded into an amyloid fiber structure, would result in a heterogeneous aggregate that is more stable, thus energetically more favorable than a non-interacting pair. In PACT, we use the Modeller^40^ software for threading a query sequence on the structure of amyloid fiber formed by Islet Amyloid Polypeptide (IAPP)^41^. To assess obtained models we proposed *ndope* score, which is a normalized version of DOPE (Discrete Optimized Protein Energy) statistical potential implemented in the Modeller software.

### PACT correctly identifies amyloid-prone peptides

In the first step, we focused on the prediction of homoaggregation, which can be considered a special case of heteroaggregation. We compared *ndope* scores obtained for models of potential homoagregates of amyloidogenic and non-amyloidogenic peptides, for which the sequences were obtained from the AmyLoad database^42^. The majority of models obtained for amyloidogenic peptides showed much lower scores (meaning more stable structures) in comparison with non-amyloids and their first quartiles of the scores were well separated (Fig.1). Differences between both groups were statistically significant. Based on the Mann–Whitney U test we were able to reject the hypothesis that the distributions of both populations were identical (*p* = 2.48*e* − 8). Considering energy difference, we built a threshold-based classifier. The classification threshold was chosen based on the Receiver Operating Characteristic (ROC) curve, as a point on the curve closest to the point (0,1), representing perfect classification (Fig. 1B). The chosen value of *ndope* score was -242. If used merely for distinguishing amyloids from non-amyloids, such a classifier was able to achieve an Area Under ROC Curve (AUC) of 0.73 and *Accuracy* of 0.77. Moreover, high values of *Sensitivity* (0.73) and *Specifictity* (0.86) were obtained. Such results are comparable with state-of-the-art amyloid predictors on the same data set (Table S1).

**Figure 1.**
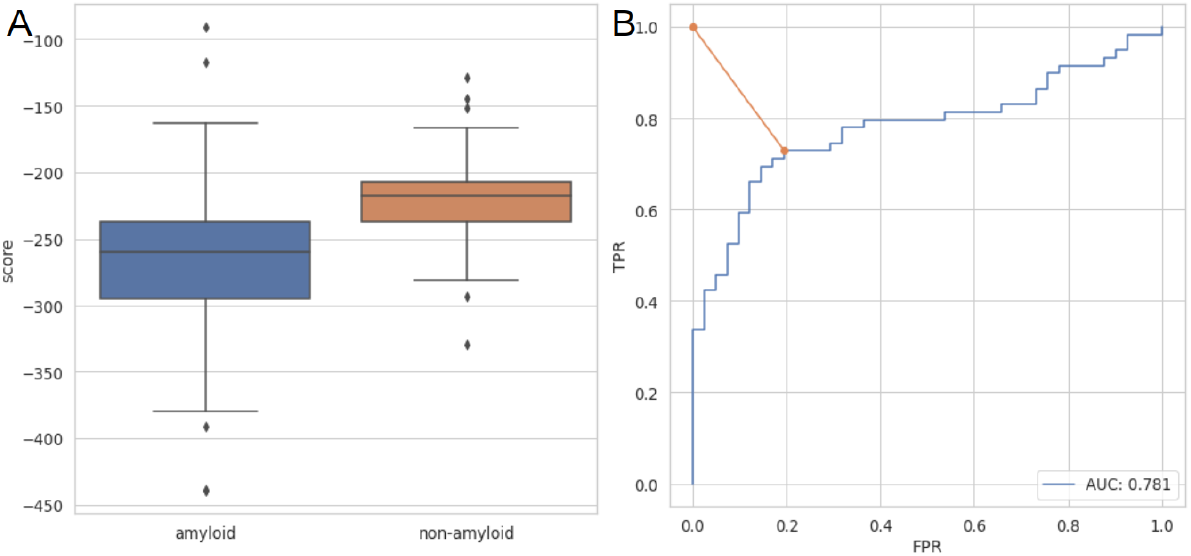
A) Distribution of *ndope* score for models of amyloidogenic and non-amyloidogenic peptides. B) ROC curve for amyloid vs non-amyloid classification. The orange line represents the distance between perfect classification point (0,1) and the chosen threshold

We also tested if the method is capable of recognizing amyloid propensity in functional amyloids, which pose a major problem for most predictors due to their under-representation in databases of amyloids. We tested the performance of the method on imperfect repeats of CsgA protein from *Escherichia coli* and *Salmonella enterica*^23^ (Fig. S1). Aggregation-prone regions of this protein (R1, R3, and R5) scored much lower than non-amyloidogenic regions (R2 and R4) from *Escherichia coli*. On this data, PACT achieved an accuracy of 0.9. Furthermore, the observed difference between *ndope* score for R4 fragments from *Escherichia coli* and *Salmonella enterica* corresponds very well to the difference in their aggregation propensities observed in experimental works^23^.

The results showed that the method can accurately predict aggregation-prone peptides of varying lengths. Furthermore, it can be utilized to detect functional amyloids.

### PACT predicts amyloid cross-interactions

We used a similar methodology to predict cross-interactions of amyloid peptides, which is the main purpose of the method. The *ndope* scores of heteroaggregates consisting of pairs of peptides whose cross-interactions resulted in faster aggregation were compared with non-amyloidogenic pairs of peptides (Fig.2). A similar analysis was performed for pairs of peptides whose cross-interactions resulted in slower aggregation (Fig. S3). In both cases, models of heterologous aggregates resulting from cross-interactions showed lower values of *ndope* scores than non-amyloids, and well-separated first quartiles of their scores (Fig.2). Furthermore, in both cases, differences between groups were statistically significant (Mann–Whitney U test, *p* = 5.54*e* − 16 for *faster* vs *negative* and *p* = 2.19*e* − 15 *slower* vs *negative* cases. Therefore, we built the threshold-based classifier using the approach described in the previous section.

**Figure 2.**
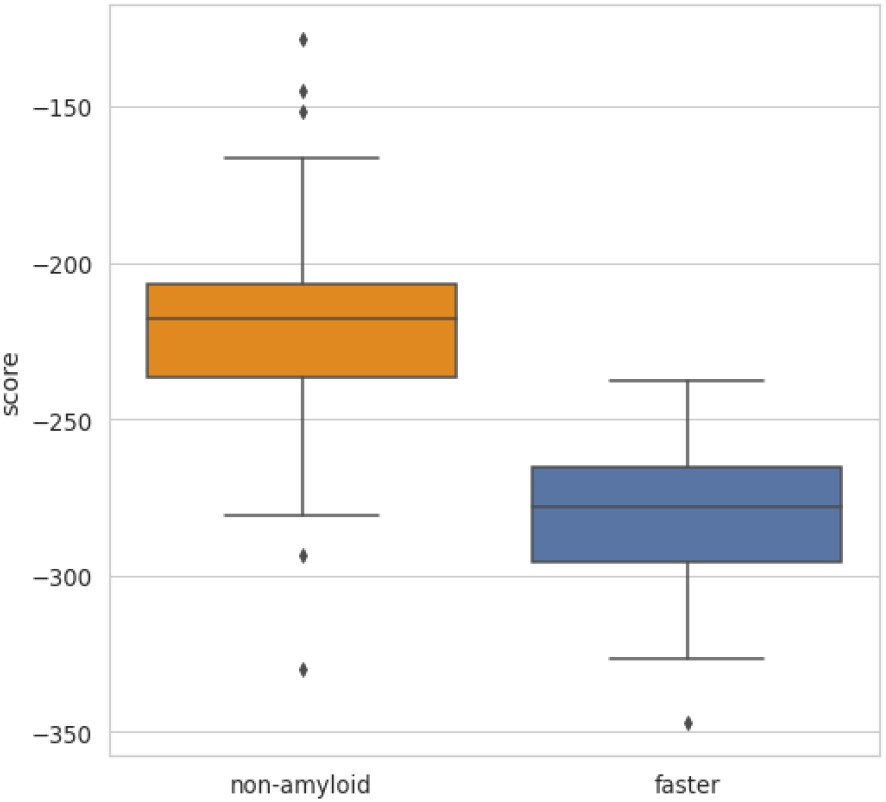
The score *ndope* for models of interacting identical non-amyloidogenic peptides (negative set) and interacting pairs resulting in increased aggregation rates (*faster* set).

To assess the performance and choose the optimal threshold value, ROC curves were calculated for both cases; *faster* rate vs *negative* (Fig.3) and *slower* vs *negative* (Fig. S3) on both training and test sets. To minimize the impact of the data choice, we performed k-folds cross-validation with k=5 on the training set and calculated several metrics describing the performance of the method (Table 1). The same metrics were then calculated on an independent test set. The same analysis was performed for the case of prediction of interactions resulting in slower aggregation (Table S2). Chosen *ndope* thresholds were very similar in both scenarios, namely -256 and -245 for faster vs negative and slower vs negative respectively. PACT performed well on both cross-validation and independent test set. It achieved the *Accuracy* of 0.83 and 0.80 on test sets of *faster* vs *negative* and *slower* vs *negative* cases, respectively. In all cases, the results obtained on the test set were within the value of one standard deviation range from the mean values obtained with the cross-validation procedures. The method performance was quite similar in both *faster* vs *negative* and *slower* vs *negative* scenarios. However, due to the smaller data set size, a larger standard deviation was obtained for *slower* vs *negative* scenario (Table 1). The results show that the method can predict whether two peptides can cross-interact but cannot distinguish between types of interactions with regard to their rate.

**Figure 3.**
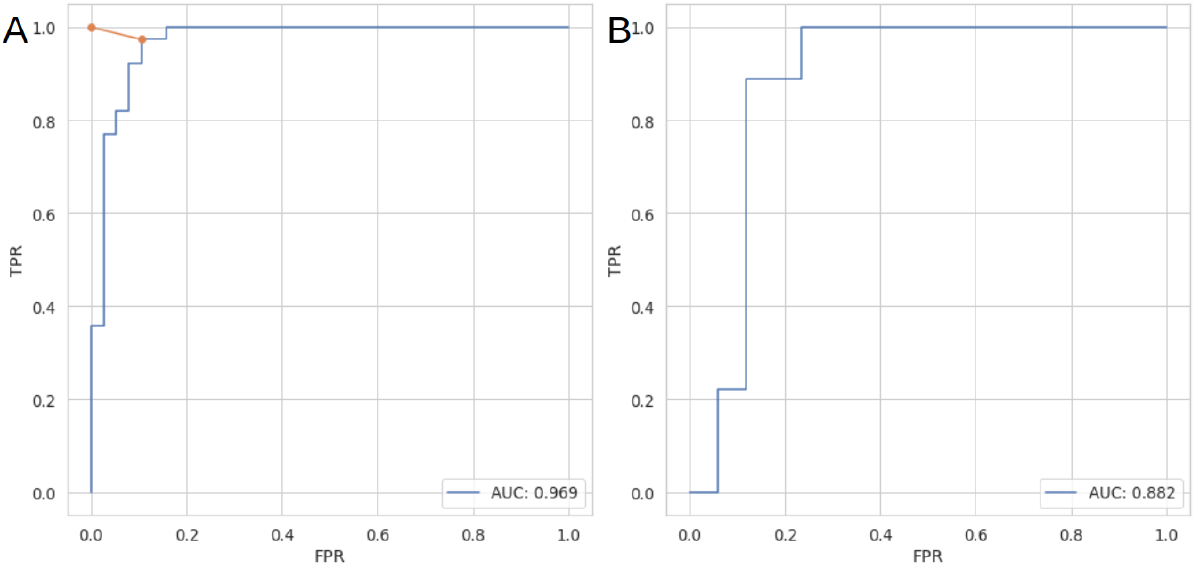
ROC curves for classification of non-aggregating and cross-interacting pairs resulting in faster aggregation on (A) training and (B) test set.

**Table 1.** Performance of PACT on cross-validation and independent test set for classification of non-aggregating and cross interacting pairs resulting in faster aggregation.

|  | Acc [std] | Sens [std] | Spec [std] | F1 [std] | MCC [std] |
| --- | --- | --- | --- | --- | --- |
| Cross-validation | 0.90 [0.05] | 0.91 [0.03] | 0.90 [0.08] | 0.90 [0.04] | 0.80 [0.06] |
| Test set | 0.83 | 0.78 | 0.88 | 0.82 | 0.66 |

### PACT is robust to bias in data

A serious problem with the data regarding interacting amyloids, which is available in the literature and, consequently, our dataset, is the large overrepresentation of interactions concerning the Abeta peptide. This may cause overfitting of the method to this group of sequences. To assess its effect, we analyzed the scores obtained for interactions between different Abeta variants (Fig. S4). The observed scores for Abeta pairs fall within the range of values observed for the remaining pairs and, therefore, they should not have a significant effect on the performance of the method. These pairs showed a relatively narrow distribution of the *ndope* values, centered slightly below the *ndope* value of −275, which is relatively close to the identified classification threshold of −256, while the remaining interacting pairs showed even lower scores.

### Interactions between bacterial amyloids and Alpha-synuclein

In recent years, numerous studies have highlighted the connection between the gut microbiome composition and the onset of many diseases, including neurodegenerative ones such as Alzheimer’s and Parkinson’s diseases^43^. Despite extensive research, understanding of the molecular mechanisms underlying this connection remains elusive. One possible explanation for this relates to functional amyloids from bacteria and human disease-related amyloids through the cross-interaction theory. The aggregation of bacterial amyloids could speed up the aggregation of disease-related proteins, leading to the disorder^44^. This hypothesis seems consistent with the results obtained by Chen and co-workers, who discovered increased production and aggregation of Alpha-synuclein in rats exposed to bacterial strains producing biofilm-related functional amyloids^45^.

In order to better understand this connection we studied possible interactions between bacterial functional amyloid CsgA and human Alpha-synuclein, whose aggregation is a hallmark of Parkinson’s disease. We used PACT to predict interactions of CsgA protein from five different organisms found in the human microbiome; *Escherichia coli* (EC), *Hafnia alvei* (HA), *Yokenella regensburgei* (YR), *Citrobacter youngae* (CY), and C*edecea davisae* (CD) with human Alpha-synuclein which were recently studied experimentally by Bhoite and coworkers^46^. It should be noted that CsgA protein was not included in the data set used to develop PACT since it exceeded the maximum length of the template.

The sequence of Alpha-synuclein was divided into overlapping fragments of length 20 amino acids and their interactions with R1-R5 repeats of each of CsgA proteins were tested. Consistently with experimental results, all of the studied CsgA variants were predicted to interact with Alpha-synuclein. Among CsgA proteins’ fragments, R1, R3 and R5 were predicted to interact, with R5 showing the best scores (Fig. 4). These results are consistent with our current state of knowledge about CsgA as the most aggregation-prone regions in these proteins are R1, R3 ad R5. Furthermore, R5 fragment which showed the lowest *ndope* scores, is located at the protein surface, therefore it can interact without a need for major conformational changes. On the Alpha-synuclein part, the best scoring region was located between positions 32 and 56 (Fig. S5). This region was recently shown to be of crucial importance for the aggregation of the protein^47,48^.

**Figure 4.**
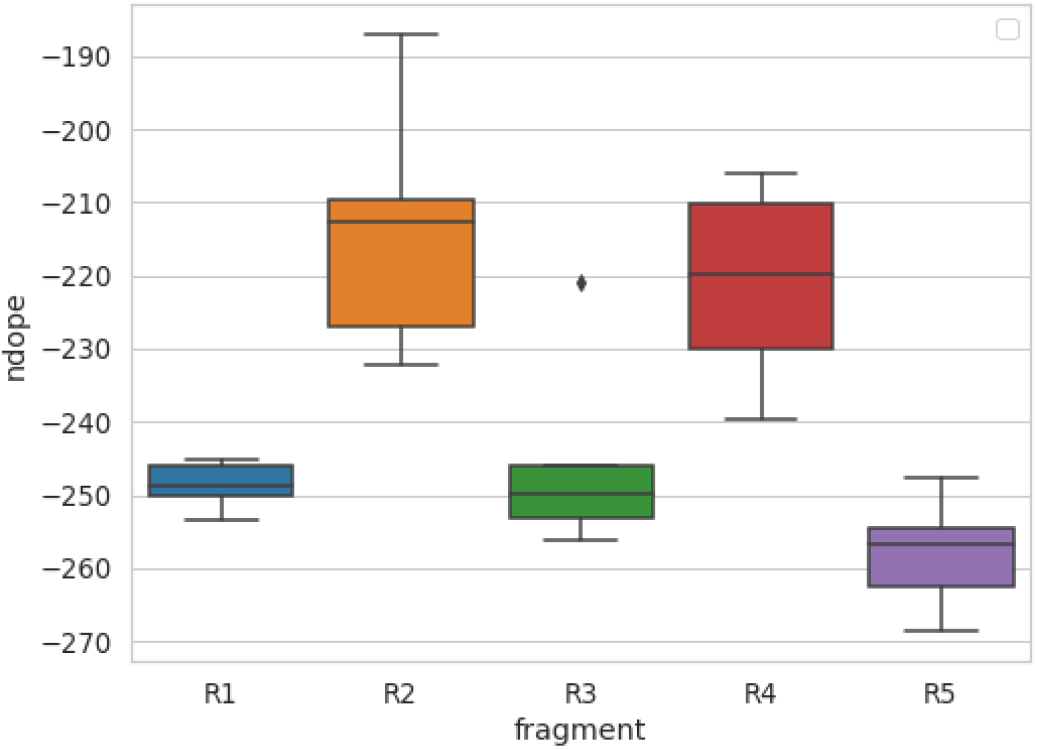
Lowest *ndope* scores for interactions of R1-R5 repeats from each of CsgA proteins with Alpha-synuclein.

### Mechanism of cross-interactions between CsgA and hIAPP

Finally, we studied interactions of CsgA protein from *Escherichia coli* with human Amylin (hIAPP) and complemented the results with experimental validation (see Supplementary materials). It was previously shown that CsgA could enhance the aggregation of hIAPP^18^. We aimed at more detailed characterization of this interaction by identifying which CsgA region is most likely to interact with hIAPP. To do so, interactions between each of CsgA repeats and hIAPP were first modeled. PACT classified positively interactions of hIAPP with R1 and R5, with the scores of -257.39 for R5 and -256.52 for R1. Notably, these fragments are likely to be exposed to the environment, which additionally makes them good candidates for potential interactions. To test the PACT predictions, experimental validation was performed using ThT assay (Methodology and results are presented in section 4 of the Supplementary materials). Obtained results showed stronger fluorescence in both cases and reduced durations of the lag phase and half-time of hIAPP aggregation in the presence of R5 (Fig. S8, Table S5). This could suggest a particular role of R5 fragment in seeding hIAPP, as predicted by PACT.

### Code Availability

PACT was implemented as an open-source Python module, available at GitHub repository: https://github.com/KubaWojciechowski/PACT. For users’ convenience, we prepared a docker container for the application, as well as the web server: https://pact.e-science.pl/pact/. For the prediction of cross-interaction we recommend the use of a default score threshold of -256 and for the prediction of homoaggregation -242. The classification result denoted as “1” indicates potential interactions. Apart from the classification, the software returns generated models of aggregates.

## Discussion

We proposed the first computational method for predicting amyloid cross-interactions. It is based on a highly interpretable and well-established physicochemical model, which is not heavily dependent on training data. This feature is especially important since the available data contains a strong interest bias towards interactions of a few popular amyloids related to neurodegenerative diseases, for example Abeta. However, in case of our method we carefully studied the effect of this overrepresentation and showed that it does not affect its performance. Furthermore, good performance on functional amyloids, which are very underrepresented in the datasets, suggests that the method is robust and can be effectively used on a wide range of sequences. In total, PACT achieved a high accuracy of 0.83 and 0.80 on the independent test sets of interactions concerning increasing and decreasing aggregation rates. On both sets the method achieved high AUC values of 0.88 and 0.89, and F1 values of 0.82 and 0.77, respectively. On the other hand, since both cases were characterized by similar interaction energies, the method cannot distinguish between enhancement and inhibition of aggregation. These results suggest that both processes may be driven by similar mechanisms. The issue was addressed in a recently published work by Louros and coworkers^49^, who applied a somewhat similar approach to study the effect of point mutations on aggregation characteristics.

We used PACT to predict the interactions of bacterial functional amyloid CsgA from different species with human Alpha-synuclein and human amylin. Although these interactions were not included in the training data set, our results are in good agreement with recently published data regarding these pairs of proteins. Importantly, they also indicate which regions can drive the cross-interactions between both proteins. The identification of potentially interacting regions can provide important insights into the possible mechanism of the process and guide future experiments.

Apart from the identification of amyloid cross-interactions, the proposed method is also capable of reliably predicting amyloid-prone regions in proteins with comparable accuracy to state-of-the-art techniques. Furthermore, it overcomes their major limitations regarding the identification of functional amyloids. Unlike most of the currently available amyloid predictors, it does not rely on the scanning of a query sequence with a very short sliding window.

High-throughput identification of amyloid cross-interactions is an important step towards our understanding of its mechanisms. It can allow for a better understanding of the principles governing the process and can also be used to identify novel cases of amyloid interactions. Such capabilities can shed light on possible mechanisms responsible for the comorbidity of devastating disorders.

## Methods

The main assumption of the method is that interactions between amyloidogenic fragments that cross-interact, threaded into an amyloid fiber structure, would result in a heterogeneous aggregate that is more stable, thus energetically more favorable than a non-interacting pair. A somewhat similar assumption was successfully applied in our previous work to predict the aggregation of short amyloidogenic fragments^36^, although the current approach differs in other aspects of the method and the objectives. In PACT, we use the Modeller software for threading a query sequence on the structure of amyloid fiber formed by Islet the Amyloid Polypeptide (IAPP)^41^. To assess obtained models we have proposed *ndope* score, which is a normalized version of the DOPE statistical potential implemented in the Modeller software.

### Data sets

To build and test the method we used the following datasets:

- the set of 86 amyloidogenic (*amyloid*) and 55 non-amyloidogenic (*non-amyloid*) peptides of lengths between 14 and 45 from the AmyLoad database^42^.
- the set of 119 pairs of peptides, which enhance (*faster* dataset) and 73 which slow down (*slower* dataset) the aggregation of each other. Both from AmyloGraph database^50^. After the removal of identical records 57 and 55 pairs of peptides which enhance and slow down the aggregation of each other respectively.

The first two sets (*amyloid* and *non-amyloid*) were used to test the method on cases of homoaggregation i.e., identifying amyloid-prone peptides.

For the prediction of cross-interactions, we used *faster, slower* and non-amyloid sets. The use of the set of non-aggregating peptides as the negative set in the interaction study was caused by the lack of a sufficient number of negative examples of non-interacting amyloid pairs. This is a common problem in studies of protein-protein interactions since negative results are rarely published, which often creates a strong bias in biological data^51^. An analysis of this dataset reveals that it is mostly composed of peptides with strong beta propensity used by authors of the Tango method^52^. The proteins from this set could be mistaken for amyloid proteins by modeling methods, therefore they provide the best available negative dataset concerning amyloidogenicity. Importantly, due to the length restrictions, CsgA protein, which was used in our validation studies, was not included in the data sets used in the development of PACT.

Datasets used in this study are available at GitHub repository: https://github.com/KubaWojciechowski/PACT

## Modeling

A query pair of sequences were threaded on the structure of amyloid fiber formed by Islet Amyloid Polypeptide (IAPP)^41^. In order to allow the method to deal with sequences of varying lengths, sequences shorter than the sequence of a template use only the main part of the template structure. In such cases, a shorter sequence is aligned to the middle of the template sequence (Fig. 5A). This choice can be justified considering that most of the currently known amyloid fragments, which are longer than a few amino acids, share a similar beta-sheet turn architecture, commonly known as the beta arch. This assumption was previously successfully applied by Ahmed and coworkers to build the ArchCandy method for amyloidogenic region prediction^53^. PACT allows sequences to be marginally longer than the template and, as a result, can be used to study cross-interactions between peptides of lengths between 14 and 45. For each of the tested pairs, 10 different models, consisting of two chains of each interacting peptide (Fig. 5B) were built using Modeller 9.24 model-multichain.py procedure with default parameters^40^. Then, the model with the lowest DOPE value was chosen for further analysis. Since the dataset consisted of fragments of varying lengths, we proposed a normalized DOPE score (*ndope*) defined as follows:

**Figure 5.**
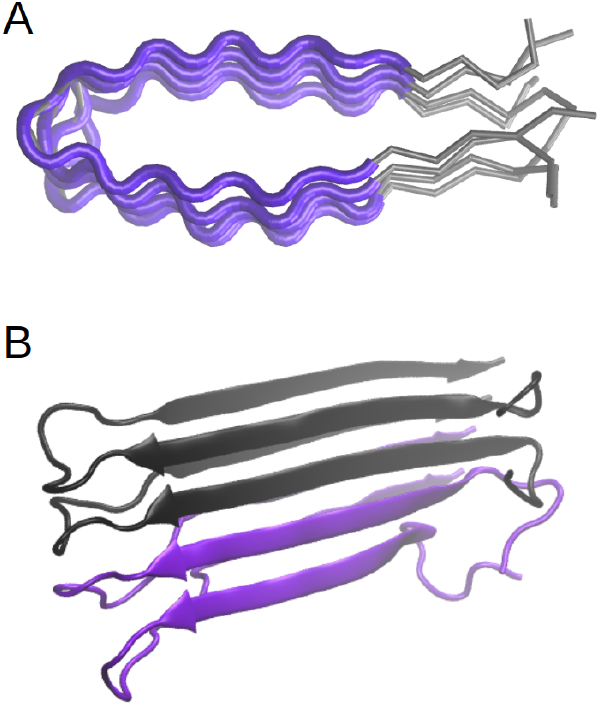
Schematic representation of the modeling procedure. A) In case when a query sequence is shorter than the template, only a part of it is used in modeling. B) The model of heterogenous fibril consists of two chains of each interacting peptide.

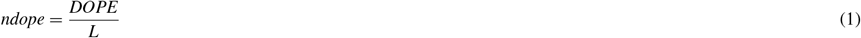

where *L* is an average length of sequences used to build a given model. Then, *ndope* scores were compared between amyloids and non-amyloids, as well as between pairs of amyloids interacting with non-amyloids.

To choose the *ndope* threshold for the classification, Receiver Operating Characteristics (ROC) curve was calculated by applying different score thresholds and recording False Positive Ratio (FPR) and True Positive Ratio (TPR). The threshold closest to the (0,1) point (representing perfect classification) was chosen. The whole procedure was schematically summarized in Fig. 6.

**Figure 6.**
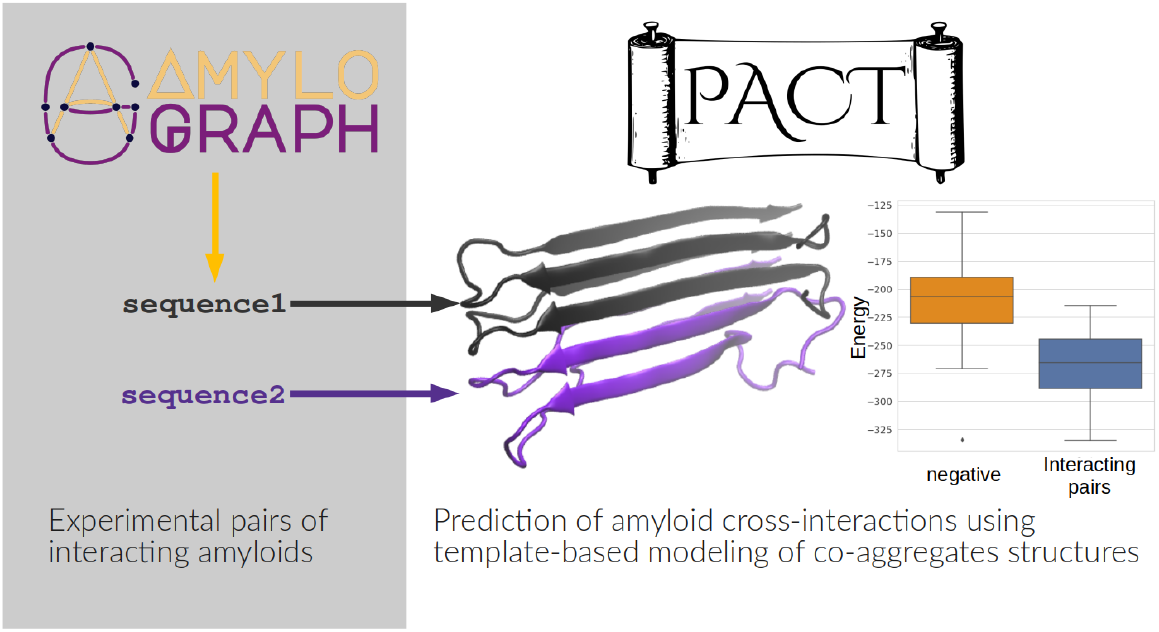
Schematic procedure of PACT

We also tested a variant of the method which utilized three different structural templates (PDB: 2nnt, 2e8d), however, it did not improve the accuracy of the method but significantly increased the computational time. Therefore, this approach was finally abandoned.

### Assessment of performance and data analysis

All the data analysis was performed using Python 3.8 with Matplotlib^54^, NumPy^55^, Pandas^56^, Scikit-Learn^57^, and Seaborn^58^ packages.

To test the performance of the proposed method, a dataset was randomly split into a training set and test set, which consisted of 30% of the data. Additionally, k-folds cross-validation (with k=5) was performed on the training data. Area Under ROC curve (AUC), *Accuracy* (ACC), *Sensitivity* (Sens), *Specificity* (Spec) and Matthew Correlation Coefficient (MCC) were used to assess the performance of the method

### Effect of Amyloid-beta variants

For the analysis of the effect of over-represented amyloid beta pairs, we divided the *faster* data set into two subsets: one containing only pairs were both interacting peptides were variants of amyloid beta (16 pairs) (*abeta*), and the set of remaining pairs (39 pairs) (*no abeta*).

### Interactions between bacterial amyloids and Alpha-synuclein

Modeling the interactions between Alpha-synuclein and CsgA proteins was performed using human Alpha-synuclein sequence (Uniprot id: P37840) and CsgA protein from five different organisms found in human microbiome; *Escherichia coli* (EC) (Uniprot id: P28307), *Hafnia alvei* (HA) (Uniprot id: G9Y7N6), *Yokenella regensburgei* (YR) (Uniprot id: A0A6H0K4L9), *Citrobacter youngae* (CY) (Uniprot id: A0A549VPM7), and C*edecea davisae* (CD) (Uniprot id: S3IYN9). The sequence of Alpha-synuclein was divided into overlaping subsequences of lengths 20. This window length was chosen because it is similar to the length of repeated units in CsgA protein, responsible for its aggregation. CsgA variants were split into five non-overlaping fragments R1-R5 corresponding to five imperfect repeats observed in their sequences. Interactions of each of CsgA fragments with all Alpha-synuclein fragments were studied.

## Data availability

Datasets used in this study are available at the GitHub repository: https://github.com/KubaWojciechowski/PACT

## Acknowledgements

We would like to thank the team of AmyloGraph developers for their effort in preparing the database used in this project and Alicja W. Nowakowska for valuable discussions of the manuscript.

This work was partially supported by the National Science Centre, Poland, Grant 2019/35/B/NZ2/03997. Access to Wroclaw Centre for Networking and Supercomputing is greatly acknowledged.

## Author contributions statement

J.W.W. and M.K. developed the concept. J.W.W. and W.S. implemented the algorithms. N.S. and M.S. performed experimental validation J.W.W. and M.K. analyzed data and wrote the manuscript.

