## Supplementary materials for "PACT - Prediction of Amyloid Cross-interaction by Threading"

#### 1. Classification of amyloid vs non-amyloid peptides

To assess PACT performance for recognizing amyloids from non-amyloids, we calculated the same metrics for three other amyloidogenicity predictors: FoldAmyloid, AmyloGram and PACT (Table S1). PACT performance was similar to these methods, which shows that it can be used also for prediction of amyloid-prone peptides.

Tablele S1 Performance of PACT on the set of aggregating and non-aggregating peptides of lengths between 14 and 45 amino acids from AmyLoad daTablease.

| Method | Accuracy | Sensitivity | Specificity | F1 | MCC |
|---|---|---|---|---|---|
| PACT | 0.77 | 0.73 | 0.85 | 0.81 | 0.55 |
| PATH | 0.67 | 0.56 | 0.85 | 0.68 | 0.41 |
| AmyloGram | 0.81 | 0.83 | 0.78 | 0.86 | 0.59 |
| FoldAmyloid | 0.75 | 0.73 | 0.78 | 0.80 | 0.49 |

#### 2. Performance on functional amyloids

Our experience shows that most of amyloidogenicity predictors perform poorly on functional amyloids, which are underrepresented in available daTableases. To test if our method can be used on functional amyloids we tested it on R1-R5 imperfect repeats from CsgA protein from *E. coli* and *S. enterica*, which we studied previously (Szulc et al. 2021). Fig.ure S2 shows calculated *ndope* scores for these fragments. Aggregation prone regions of R1, R3, and R5 scored much lower than non-amyloidogenic regions, R2 and R4, from *E. coli*.  On this data, PACT achieved the Accuracy of 0.9. Furthermore, the observed a difference in *ndope* score for R4 fragments from *E. coli* and *S. enterica, which* corresponds very well to the difference in their aggregation propensity.

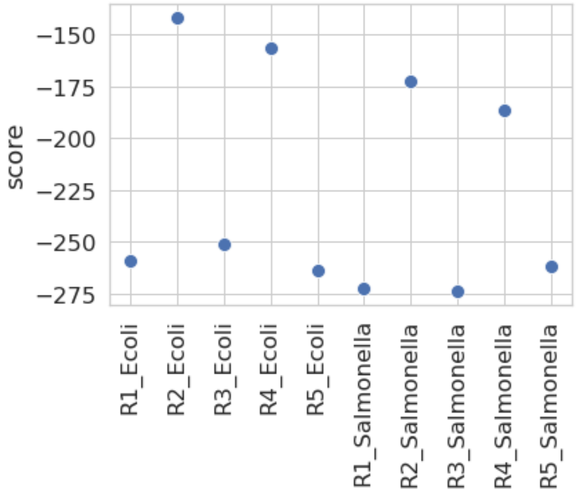


Fig. S1 ndope scores for R1-R5 imperfect repeats of CsgA protein from *E. coli* and *S. enterica.*

#### 3. Prediction of cross-interactions

In the next step, we tested the performance of the method on pairs of interacting amyloids. We tested pairs whose interactions resulted both in increased and decreased aggregation speed. The first case is described in more detail in the main text. Here we show the results for the case of interactions resulting in slower aggregation (Fig. S2).

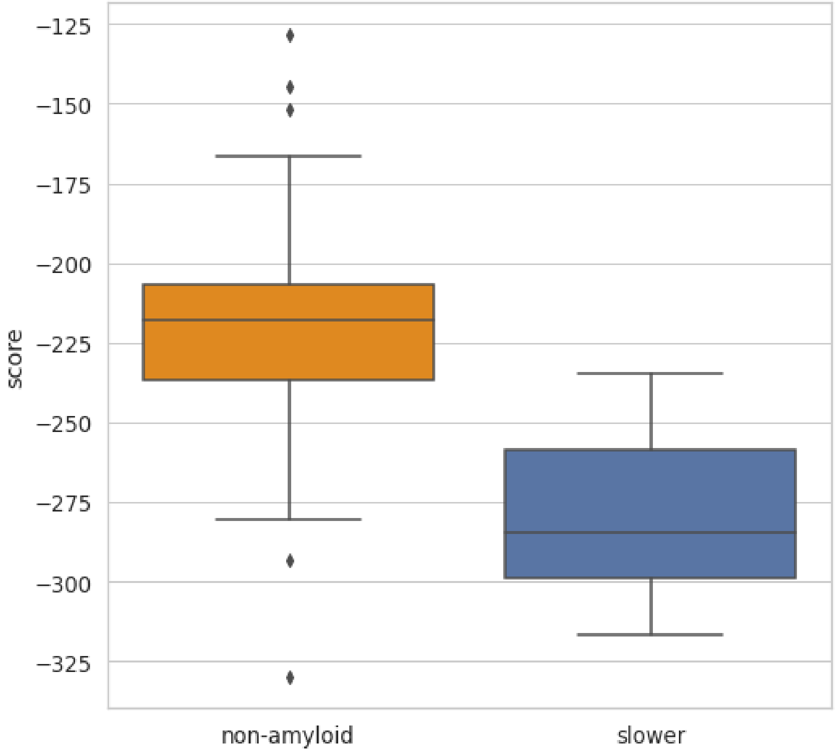

Fig. S2 Normalized DOPE score for models of non-amyloidogenic peptides and pairs of interactions resulting in decreased aggregation rates.

Same as in the case of prediction of homoaggregation, the threshold based classifier was built, but this time the data set was first split into training and test sets. On the training set k-folds cross-validation was performed with k=5. Then we used the whole training set to find the threshold again and tested the method on the independent test set. ROC curves were calculated for both training and test sets (Fig. S3), *ndope* threshold of -245 was found and other metrics were calculated (Table S2). The results similar to those obtained for faster aggregation were obtained.

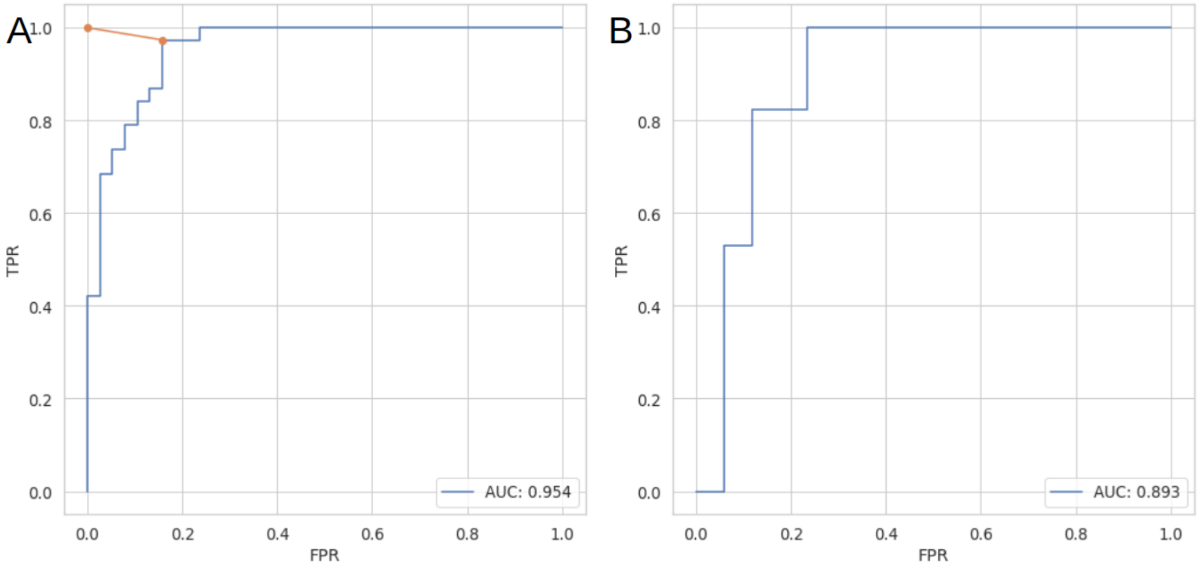


Fig. S3 ROC curves for classification of non-aggregating and cross interacting pairs resulting in slower aggregation on (A) training and (B) test set.

Table S2 Performance of PACT on cross-validation and independent test set for classification of non-aggregating and cross-interacting pairs resulting in their slower aggregation.

| | Accuracy [std] | Sensitivity [std] | Specificity [std] | F1 [std] | MCC [std] |
|---|---|---|---|---|---|
| Cross-validation | 0.90 [0.07] | 0.91 [0.17] | 0.86 [0.13] | 0.89 [0.09] | 0.81 [0.12] |
| Test set | 0.79 | 0.71 | 0.88 | 0.77 | 0.59 |

Next, we tested how interest bias of authors of the publications influences the performance of PACT. To do so, highly overrepresented interactions of different variants of Abeta were closely studied. The obtained scores for *A*beta pairs were within the same range as values for the remaining pairs, and therefore should not have a significant effect on the performance of the method. These pairs obtained quite similar *ndope* values, centered slightly below *ndope* value of -275, which is relatively close to identified classification threshold for *faster vs negative* scenario of -256.

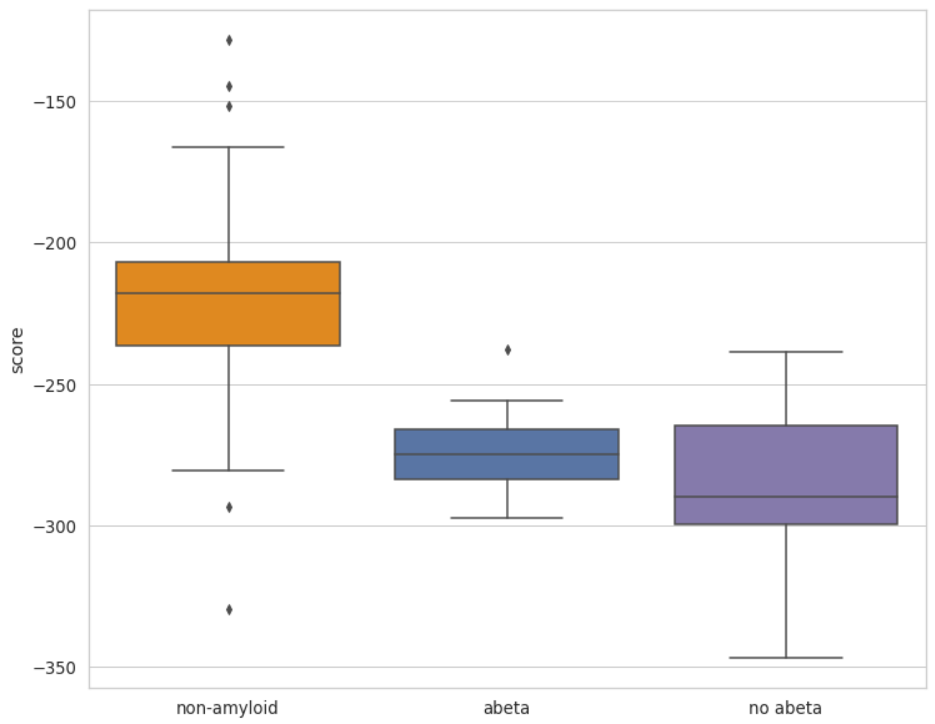


Fig. S4 Normalized DOPE score for models of non-amyloidogenic peptides and pairs of interactions resulting in increased aggregation rates where both partners belong to Abeta variants and the remaining pairs.

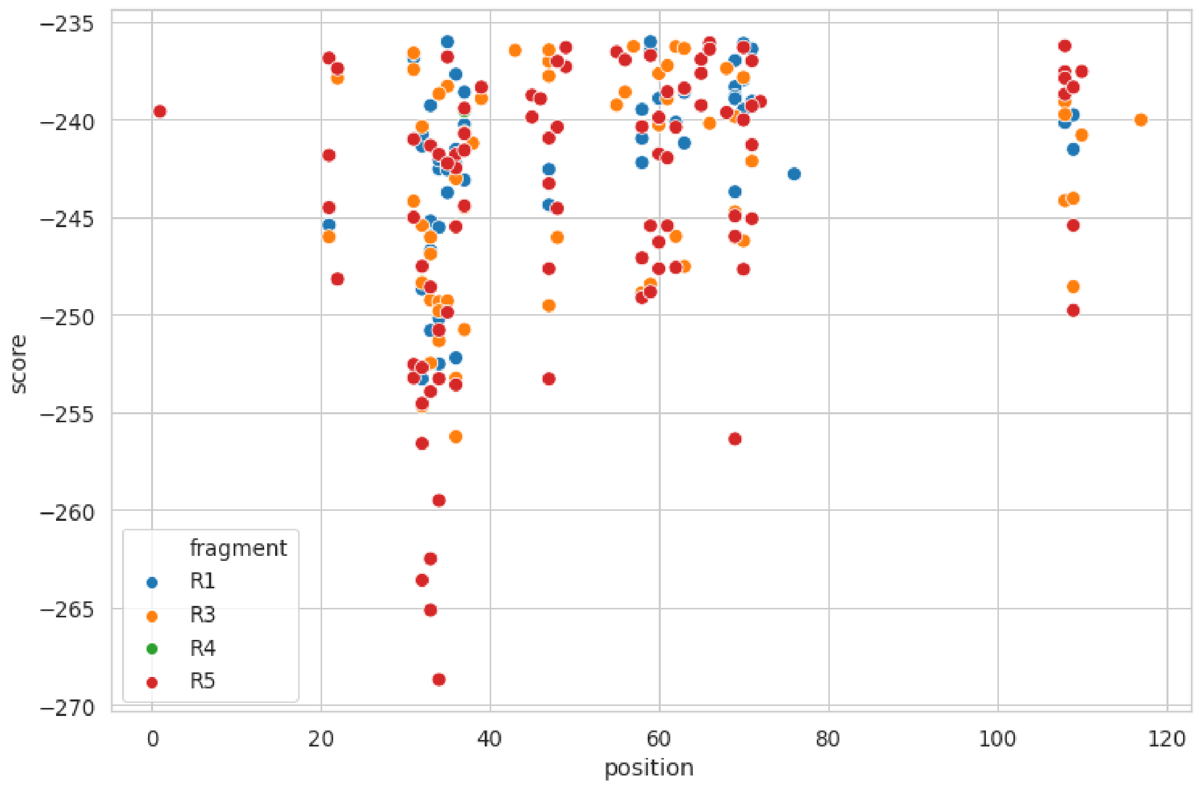


Fig. S5 Scores for interactions of CsgA fragments with Alpha-synuclein fragments. Each dot represents the starting position of 20 amino acid fragment of the sequence.

### 4. Experimental validation

Table S3 Peptides analytical data.

| Name | Sequence | Formula | Calculated M/z | Experimental M/z | Analytical HPLC $t_r$ [min] |
|---|---|---|---|---|---|
| R1 | H-SELNIYQYGGGNSALALQTDARN-$NH_2$ | $C_{94}H_{153}N_{29}O_{36}$ | [(M+2H)/2] 1228.1060<br>[(M+3H)/3] 819.0733 | [(M+2H)/2] 1228.1328<br>[(M+3H)/3] 819.0726 | 14.599 |
| R5 | H-SDLTITQHGGGNGADVGQGSDD-$NH_2$ | $C_{87}H_{138}N_{28}O_{36}$ | [(M+2H)/2] 1994.9397<br>[(M+3H)/3] 997.9738 | [(M+2H)/2] 1994.2070<br>[(M+3H)/3] 997.5491 | 12.273 |
| hIAPP | H-KCNTATCATQRLANFLVHSSNNFGAILSSTNVGSNTY-$NH_2$ | $C_{165}H_{261}N_{51}O_{55}S_2$ | [(M+3H)/3] 1302.8<br>[(M+4H)/4] 977.3 | [(M+3H)/3] 1302.5<br>[(M+4H)/4] 977.3 | 10.230 |

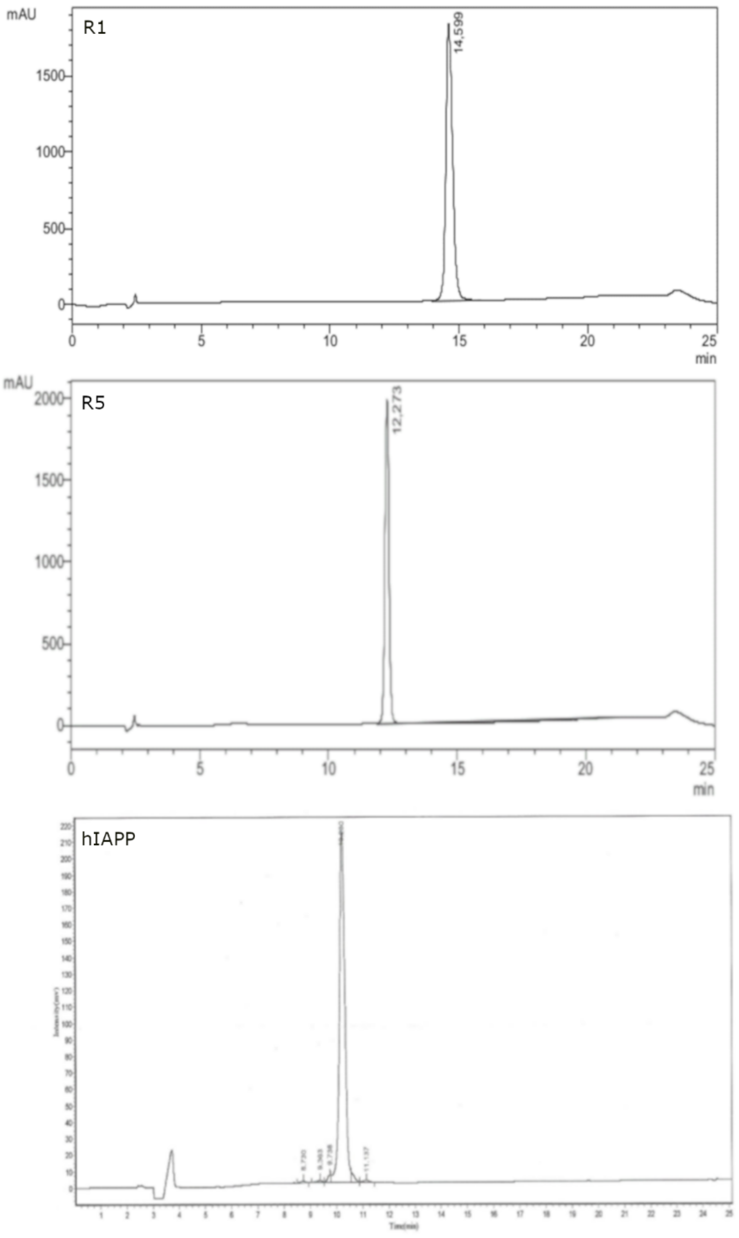


Fig. S6 Analytical HPLC chromatograms of the studied peptides.

#### Materials and Methods

**Peptide synthesis.** All commercially available reagents and solvents were purchased from Merck and used without further purification. Peptides R1 and R5 were synthesized with an automated solid-phase peptide synthesizer (Liberty Blue, CEM) using H-Rink amide ChemMatrix resin 35-100 mesh particle size (loading: 0.59 mmol/g). Fmoc deprotection was obtained using 20% piperidine in DMF for 1 min at 90 ºC. A single-coupling procedure was achieved with 0.5 M solution of *N,N′*-diisopropylcarbodiimide (DIC) and 0.5 M solution of Oxyma Pure Novabiochem® in DMF for 4 min at 90 °C. Cleavage of the peptides from the resin was accomplished with the mixture of TFA/TIS/$H_2O$ (95:2.5:2.5) after 3 h of shaking. The crude peptide was precipitated with ice-cold $Et_2O$ and centrifuged (7 000 rpm, 10 min, 4 °C). Peptides were purified using preparative HPLC (Knauer AZURA ASM 2.1L) with a C18 column (Thermo Scientific, Hypersil Gold 12 µm, 250 mm × 20 mm) with water/acetonitrile (0.05% TFA) eluent system. hIAPP was purchased from ProteoGenix, see Table S3.

**Analytical high-performance liquid chromatography (HPLC).** Analytical HPLC for R1 and R5 was performed using column ReproSil Saphir C18 100Å 5µ 4.6 × 150 mm; detection wavelength 222 nm; eluent system: A = $H_2O$+0.05% TFA, B = $CH_3CN$+0.05% TFA, gradient: t=0–20 min, 90%–0% A; t=20–22 min, 0% A; t=22–25 min, 0%–90% A, see Fig. S6). Analytical HPLC for hIAPP was provided by ProteoGenix and performed on PLRP-S column 100Å 4.6 × 250 mm; detection wavelength 220 nm; eluent system: A = $CH_3CN$+0.1 % TFA, B = $H_2O$+0.1% TFA, gradient: t=0–25 min, 10%–90% A; t=25–30 min, 100%–0% A).

**Mass spectrometry (MS).** Peptides R1 and R5 were studied by WATERS LCT Premier XE System consisting of a high resolution mass spectrometer with a time of flight (TOF) using electrospray ionization (ESI). MS analysis for hIAPP was provided by ProteoGenix.

**Circular dichroism (CD).** CD spectra were recorded on JASCO J-1500 at 20 °C between 250 and 190 nm in water with following parameters: 0.2 nm resolution, 1.0 nm band width, 20 mdeg sensitivity, 0.25 s response, 50 nm/min scanning speed, 5 scans, 0.1 cm cuvette path length. The CD spectra of the 10 mM PBS buffer pH 7.4 was recorded and subtracted from the raw data. The peptides were dissolved in hexafluoroisopropanol (HFIP), then mixed for 3 hours to obtain monomers. HFIP was evaporated overnight in a desiccator, then the samples were dissolved in a PBS buffer to obtain peptide concentration of 100 µM. Then, a filtration process was conducted, and the resulting filtrate was employed for subsequent experimentation. The CD intensity is given as mean residue ellipticity (θ [deg × $cm^2$ × $dmol^{-1}$]). The spectra were smoothed using the Savitzky–Golay filter (polynomial order 2, widow size 19) applied in the SciPy package.

**Thioflavin T (ThT) fluorescence assay.** Kinetic measurements were carried out in a 96-well BRANDplate® on a CLARIOstar Plus, BMG LABTECH, at 20 °C, using wavelengths of 440±15 nm and 480±20 nm, for ThT excitation and emission respectively. Additionally, the plate was shaken for 30 s at the interval of 30 min during 24 hours of measurements. Final concentrations were 50 µM of ThT and 100 µM of each monomerized peptide. Peptides were monomerized according to the procedure described in the CD section. The experiment was performed in the duplicate. The obtained fluorescence values were normalized to the fluorescence maximum in the 0–1 range.

#### Results

An experimental validation was conducted to confirm the predictions of cross-interactions obtained by PACT. Peptides for the studies were chosen based on the predicted energies and availability and included fragments R1 and R5 of the CsgA protein from *E. coli* species, known for their functional amyloid properties[1], as well as hIAPP, an amyloid associated type 2 diabetes[2]. The interactions between the N-terminal (R1) and C-terminal (R5) fragments of CsgA protein and hIAPP were investigated using circular dichroism (CD) and Thioflavin T (ThT) fluorescence assays. The studies were undertaken to demonstrate cross-interaction of R1 and R5 fragments with hIAPP, as predicted computationally with PACT.

CD spectra of all samples exhibited a minimum at approximately 200 nm upon dissolution (see Fig. S7 and Table S4) indicating a random coil formation. Throughout the time course of the experiment, only a slight shift towards lower wavenumbers (approximately 1-2 nm) was observed for all samples. Despite that the secondary structure characteristic observed still resembled those of a random coil, the reduced spectral intensity and higher voltage on the photomultiplier indicate occurrence of the temporal aggregation.

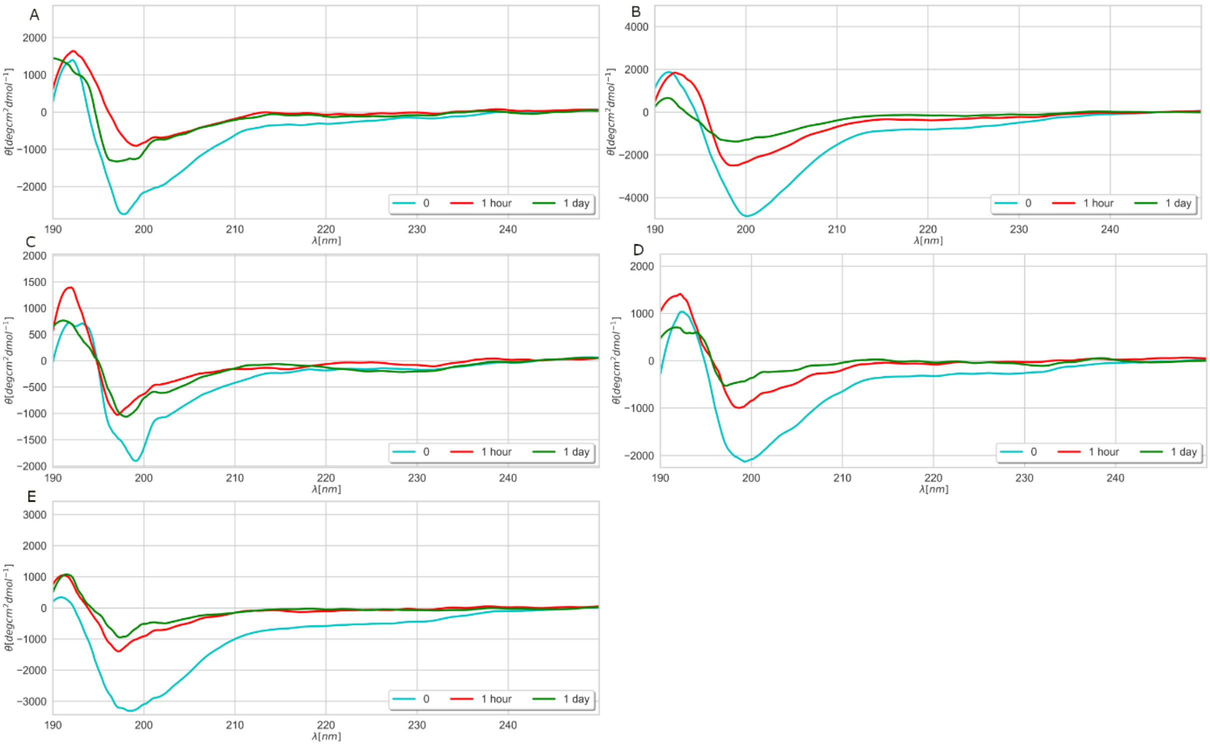


Fig. S7 Far-UV CD spectra of the studied samples: (A) fragment R1 of CsgA protein of *E. coli* species, (B) fragment R1 of CsgA protein of *E. coli* species + hIAPP, (C) fragment R5 of CsgA protein of *E. coli* species, (D) fragment R5 of CsgA protein of *E. coli* species + hIAPP, (E) hIAPP, on the day of dissolving, after one hour and after one day in the PBS buffer. Cpep=100 µM.

Table S4 Changes in the positions of CD spectra minima (given in nm) of the studied samples within time in the PBS buffer. Cpep=100 µM.

| sample / time | R1 | R5 | hIAPP | R1+hIAPP | R5+hIAPP |
|---|---|---|---|---|---|
| 0 | 197.8 | 199.2 | 198.6 | 200 | 199.2 |
| 1 hour | 199.2 | 197.2 | 197.2 | 198.6 | 198.6 |
| 1 day | 197.0 | 198.0 | 197.4 | 198.2 | 197.2 |

Comparative analysis revealed an acceleration of aggregation for the R1+hIAPP and R5+hIAPP samples compared to the individual peptides (see Fig. S8 and Table S5), confirming cross-interactions between the peptides. This acceleration was accompanied by a reduction in the lag phase duration from hours, observed in the case of terminal fragments of the CsgA protein, to minutes upon addition of hIAPP to the mixture. Additionally, the heterogeneous mixture of R5+hIAPP showed a faster rate of aggregation compared to R1+hIAPP (see Table S5). The R5+hIAPP mixture exhibited a stronger cross-interaction effect than R1+hIAPP. The PACT score indicated a lower energy (-257.39) for the R5-hIAPP interaction compared to R1+hIAPP (-256.52), which is consistent with the ThT results. The difference between these two heterogeneous mixtures was minimal based on the half-time, lag time, and the increase in normalized fluorescence value.

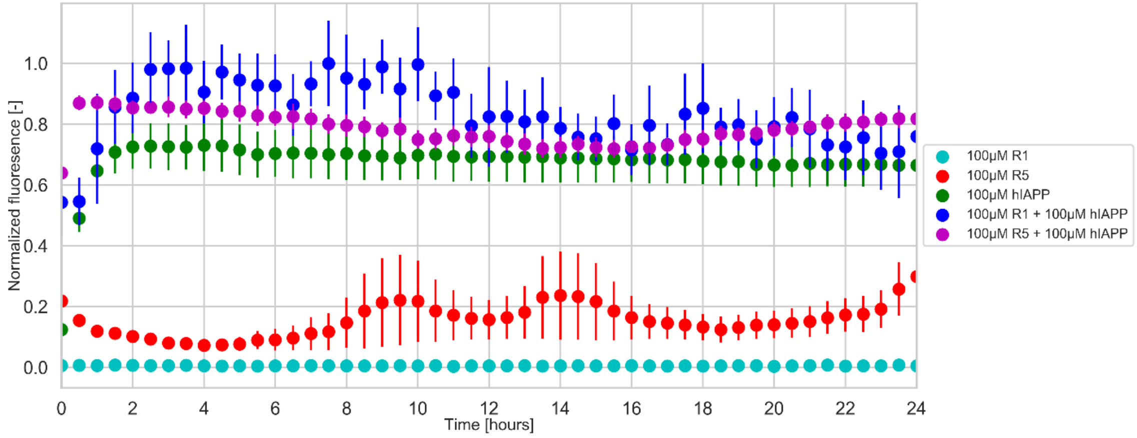


Fig. S8 ThT curves for the studied samples following the aggregation process in the PBS buffer. Cpep=100 µM.

Table S5 Parameters obtained by fitting aggregation kinetics to studied peptides. Cpep=100 µM. Where, the symbol '-' denotes that no half-time calculation was performed due to the absence of a fitted curve (only the lag phase was observed, representing the monomeric state of the peptide).

| sample / parameter | R1 | R5 | hIAPP | R1+hIAPP | R5+hIAPP |
|---|---|---|---|---|---|
| lag phase | 24 hours | 23 hours | 30 minutes | 30 minutes | 10 minutes |
| half time | - | 24 hours | 50 minutes | 1 hour | 30 minutes |
